# Biomechanical stress analysis of Type-A aortic dissection at pre-dissection, post-dissection, and post-repair states

**DOI:** 10.1101/2023.10.06.561307

**Authors:** Christina Sun, Tongran Qin, Asanish Kalyanasundaram, John Elefteriades, Wei Sun, Liang Liang

## Abstract

Acute type A aortic dissection remains a deadly and elusive condition, with risk factors such as hypertension, bicuspid aortic valves, and genetic predispositions. As existing guidelines for surgical intervention based solely on aneurysm diameter face scrutiny, there is a growing need to consider other predictors and parameters, including wall stress, in assessing dissection risk. Through our research, we aim to elucidate the biomechanical underpinnings of aortic dissection and provide valuable insights into its prediction and prevention.

We applied finite element analysis (FEA) to assess stress distribution on a rare dataset comprising computed tomography (CT) images obtained from eight patients at three stages of aortic dissection: pre-dissection (preD), post-dissection (postD), and post-repair (postR). Our findings reveal significant increases in both mean and peak aortic wall stresses during the transition from the preD state to the postD state, reflecting the mechanical impact of dissection. Surgical repair effectively restores aortic wall diameter to pre-dissection levels, documenting its effectiveness in mitigating further complications. Furthermore, we identified stress concentration regions within the aortic wall that closely correlated with observed dissection borders, offering insights into high-risk areas.

This study demonstrates the importance of considering biomechanical factors when assessing aortic dissection risk. Despite some limitations, such as uniform wall thickness assumptions and the absence of dynamic blood flow considerations, our patient-specific FEA approach provides valuable mechanistic insights into aortic dissection. These findings hold promise for improving predictive models and informing clinical decisions to enhance patient care.

## 1. INTRODUCTION

Type A aortic dissection, a rare but catastrophic cardiovascular event, poses a significant clinical challenge due to its high mortality and morbidity rate [1]. It is initiated by the development of a tear in the intima of the aorta, which allows pressurized blood to enter the aortic wall, subsequently leading to the separation of aortic wall layers. This condition can compromise branch vessel inflow, cause true lumen compression, and result in severe complications such as acute aortic valve regurgitation, myocardial infarction, stroke, visceral organ ischemia, or frank aortic rupture and death [1, 2].

Dissections of the ascending aorta (Type A) are both more common and significantly more dangerous than descending aortic dissections (Type B). Approximately 60% of aortic dissections are Type A [1, 3]. If left untreated, the mortality rate of thoracic aortic aneurysm and dissection (TAAD) patients increases 1-2% every hour for the first 48 hours and up to 68% within the first two days [4, 5], whereas the overall in-hospital mortality rate of Type B aortic dissection patients is a much lower 11% in the first month [1, 6]. While certain risk factors, such as hypertension, bicuspid aortic valve, and genetic predispositions (e.g., Marfan syndrome) have been identified [2, 7, 8], the precise triggers of TAAD initiation remain elusive.

The traditionally employed American and European guideline for mitigating the risk of aortic dissection is the prophylactic surgical replacement of the ascending aorta when an aneurysm reaches a diameter of 5.5 cm [9, 10]. However, large retrospective studies have cast doubt on the effectiveness of this guideline, suggesting that it may be an inadequate predictor [11, 12]. Very recently, both the American and European societies have decreased the recommended general threshold for intervention to 5.0 cm [13–16]. From an engineering perspective, aortic dissection can be viewed as a biomechanical failure of the aortic wall, occurring when wall stress exceeds wall strength [15, 16].

Computational modeling has emerged as a valuable tool for studying ascending aortic aneurysms, yet the current computational studies have not been able to precisely determine when and where the dissection would occur, limiting their predictive power for clinical use. To understand the biomechanical aspects of aortic dissection, studies have explored factors associated with wall shear stress and wall principal stress, and their impact on the aortic wall structural integrity. Bäumler et al. presented a computational framework for simulating aortic dissection, considering factors of vessel wall deformation, blood flow, and pressure in patient-specific models, concluding that the flexibility of the dissection membrane significantly affects local blood flow patterns [17]. A study by Martufi et al. quantitatively characterized the geometry of abdominal aortic aneurysms using CT data, proposing that various geometric indices such as size, shape, and curvature-based metrics may be necessary to evaluate the risk of rupture [18]. He et al. utilized machine learning to predict the local strength of ascending thoracic aortic aneurysm tissues based on tension-strain data, suggesting that early phase response features can reliably estimate tissue strength [19]. Research by Nathan et al. explored the relationship between regional wall stress and aortic dissections, notably near the sinotubular junction and the left subclavian artery, suggesting a potential role in the pathogenesis of aortic dissections [20].

One limitation of previous studies is the lack of complete longitudinal data sets that cover a patient’s pre-dissection state, post-dissection state, and post-surgical repair state, which are usually difficult to obtain from clinics. The greatest issue is that it is extremely uncommon for acute dissection patients to have undergone CT imaging in close temporal proximity to the date of occurrence of the acute ascending dissection. Moreover, engineering analysis of these patient-specific aortic wall mechanics often requires the patient-specific material properties of these patients.

This research paper aims to contribute to the understanding of aortic dissection by investigating the change in aortic stress distributions at different time points in the aortic disease. By assessing aortic stress distributions using noninvasive imaging techniques and finite element analysis (FEA), we seek to reveal increased stresses in patients who have sustained acute type A aortic dissection. Importantly, we possess a rare dataset that includes pre-dissection (preD), post-dissection (postD), and post-repair (postR) data from the same patients, allowing for analyses across these time points. This comprehensive dataset sets our research apart from existing studies in the field.

## 2. METHODS

### 2.1 Patient clinical data

As seen in Table 1, clinical data from eight patients between the age of 41 and 75 were collected retrospectively from Yale New Haven Hospital. An engineering analysis was conducted on this rare dataset comprised of retrospectively obtained computed tomography (CT) images. The dataset included pre-dissection (PreD) and post-dissection (PostD) CT images from the eight patients. Four of these eight patients have an additional CT scan after surgical repair (postR). The CT images were acquired at the early systolic phase [21, 22]. The PreD CT scans were originally performed as part of the patients’ regular medical care, under the direction of their treating physicians, and were not specifically obtained for the purpose of this study.

**Table 1:** Patient Information.

| Patient ID | 1 | 2 | 3 | 4 | 5 | 6 | 7 | 8 |
| --- | --- | --- | --- | --- | --- | --- | --- | --- |
| Age at first presentation | 67 | 54 | 66 | 63 | 73 | 41 | 63 | 75 |
| Sex | M | M | F | M | F | M | M | M |
| Height (cm) | 163 | 170 | 163 | 163 | 160 | 175 | 188 | 188 |
| Weight (kg) | 120.5 | 80 | 55 | 49 | 62 | 72 | 100 | 109 |
| Max Asc Diam (cm) | 5.2 | 6 | 4.5 | 5.38 | 3.36 | 5.1 | 6.5 | 5.1 |
| PreD CT Date (mm/dd/yy) | 8/8/19 | 1/27/19 | 6/18/18 | 10/1/13 | 7/4/14 | 1/4/12 | 6/24/09 | 3/15/10 |
| PostD CT Date (mm/dd/yy) | 12/12/19 | 8/6/19 | 6/20/18 | 1/4/16 | 7/8/15 | 6/20/14 | 4/24/12 | 1/25/11 |
| PostR CT Date (mm/dd/yy) | 2/6/20 | 8/17/19 | N/A | N/A | N/A | N/A | 12/5/13 | 2/14/11 |

Note: “Max Asc Diam” refers to the maximum diameter of the ascending aorta.

### 2.2 Patient-specific geometry reconstruction and meshing

The patient-specific geometries were segmented from CT images using 3D Slicer software [23] (Figure 1). The reconstructed geometries (Figure 2A) include the aortic root, ascending aorta, aortic arch with its 3 branches (brachiocephalic artery, left common carotid artery, and left subclavian artery), the proximal and distal descending aorta, as well as the true lumen and false lumen. Finite element (FE) meshes were then generated using HyperMesh (Altair Engineering, Inc., MI) software. First, the surface geometries were generated based on the segmented patient-specific geometries (Figure 2B). Then, surface meshes with 4-node quadrilateral shell elements were created (Figure 2C). Aortic wall solid meshes with four layers of 8-node brick elements were created by offsetting the surface meshes (Figure 2D). In this study, the thickness of the aortic wall without dissection was assumed to be 2 mm, while the thickness of the dissection wall was assumed to be half of the normal thickness according to the observation of Dr. Elefteriades in clinics. The replacement graft is assumed to have a thickness of 0.3mm based on the experimental data [24].

**Figure 1:**
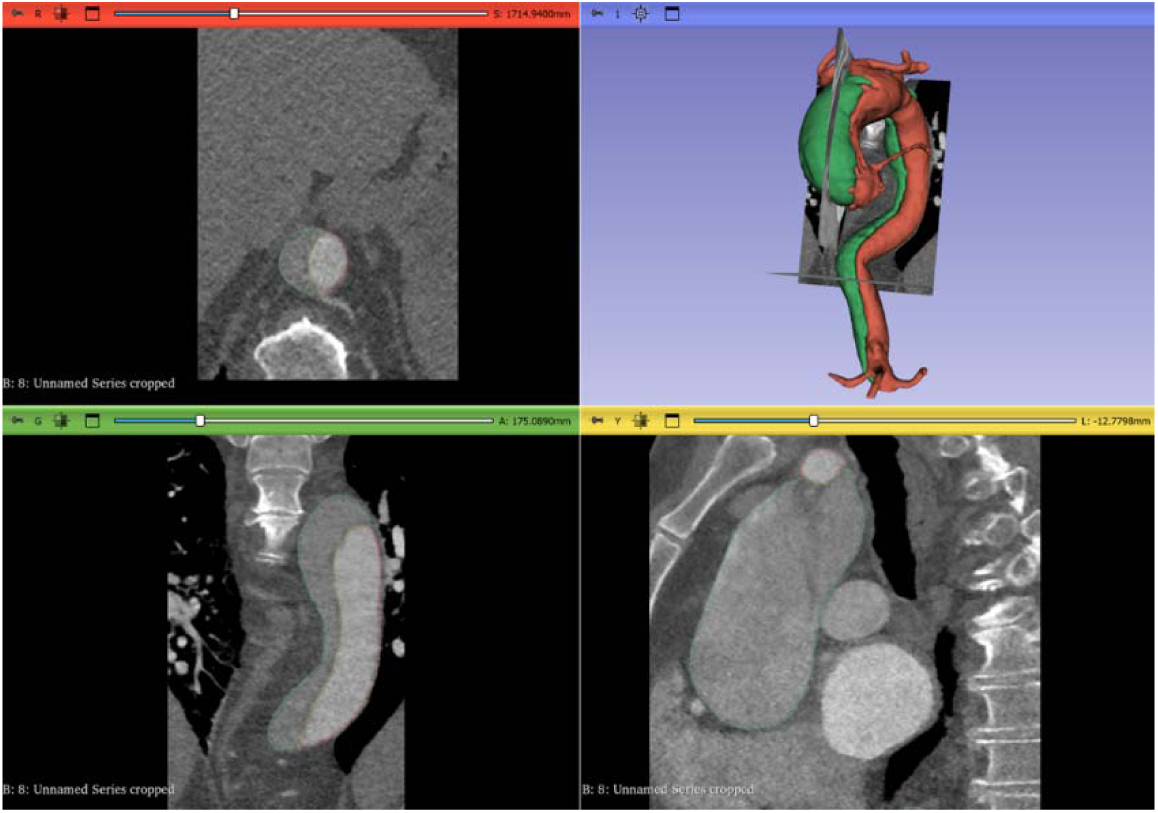
Reconstruction of patient-specific geometries based on CT data.

**Figure 2:**
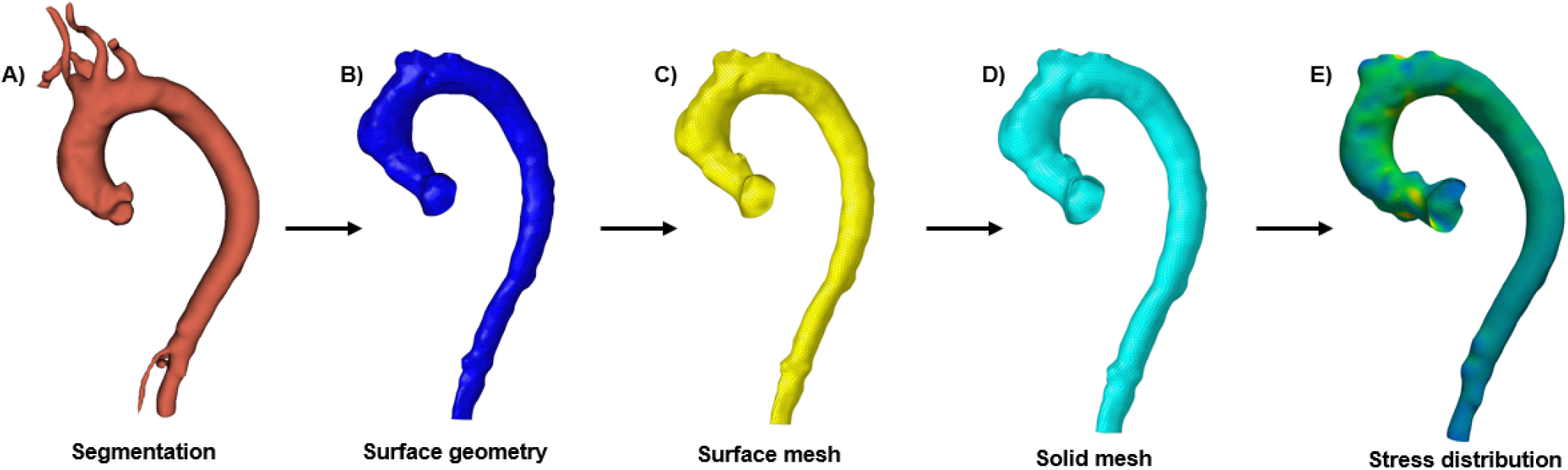
The procedure of stress analysis based on patient-specific geometries. A) a geometric model is segmented from CT images using 3D slicer software, B) surface editing using Hypermesh software, C) surface remeshing using Hypermesh, D) solid element mesh, E) FE stress analysis result.

### 2.3 Finite element (FE) modeling for stress analysis

Finite Element (FE) modeling was performed using the Abaqus/Standard (SIMULIA, Providence, RI), based on the Static Determinacy Approach (SDA). More details about the application of the SDA can be found in our previous publications [25, 26]. Briefly, static determinacy is a property of pressurized structures whose internal tension/stress is independent of the material properties (i.e., the tension/stress can be calculated based only on the external pressure/force and geometry) [26]. Using Laplace’s law to compute the wall hoop stress of a perfect cylindrical tube is one of the most well-known examples of static determinacy [26]. Though the geometry of the aorta is much more complex, it has been shown [26–28] that the aorta under a specific pressure loading can also be treated as a structure with static determinacy, and that the aortic wall stress is independent of material properties [29–31] and can be calculated directly from the pressure and geometry. Therefore, the calculation of aortic wall stress does not need material properties, and the FE modeling was done by following previous studies [26–28]. The nodal displacements were fixed at the boundaries of the aorta segments of interest, including the inlet of the ascending aorta (around the aortic annulus), the outlet of the three arch branches, and the outlet of the descending aorta. The systolic blood pressure was applied to the inner surface of the aorta, and the stress fields were calculated and analyzed (Figure E).

FE stress analysis was performed for each patient at pre-dissection (PreD) and post-dissection (PostD) states, as well as post-repair (PostR) state if data were available. To understand the impact of varying blood pressure levels on the aortic wall stresses, three systolic blood pressure values were utilized: 16 kPa (∼120 mmHg), 18 kPa (∼135 mmHg), and 20 kPa (∼150 mmHg), which corresponded to normal systolic blood pressure, hypertension stage 1, and hypertension stage 2, respectively [32].

Principal stresses were obtained for each element in the FE mesh, and the one with the maximum absolute value (a.k.a. maximum absolute principal stress) was used in stress analysis throughout this study and was simply referred to as “stress”. Because the aortic wall is meshed with 4 layers of elements across the wall thickness, the aortic wall stress is calculated as the averaged stress of the 4 layers of elements across the wall thickness. The aorta was segmented into five anatomic segments, i.e., the aortic root, ascending aorta, aortic arch, proximal descending aorta, and distal descending aorta (Figure 3). As clinical dissections occur near the root area, the aortic root was further split into the non-coronary Sinus of Valsalva (NCS), left coronary sinus (LCS), and right coronary sinus (RCS), as shown in Figure 4. The segmentation of the aorta into different regions follows the previous work [33, 34]. The mean stresses of all elements within each of the five anatomic segments of the aorta and each of the three segments of the aortic root were obtained, as a high mean stress would indicate that the entire segment is subject to high stress. The peak stresses, or the maximum stresses, of all elements within each of the five anatomic segments of the aorta and each of the three segments of the aortic root, were also obtained and compared, as a high peak stress could indicate a concentrated “hotspot” of high stress, leading to a focal location of material failure.

**Figure 3:**
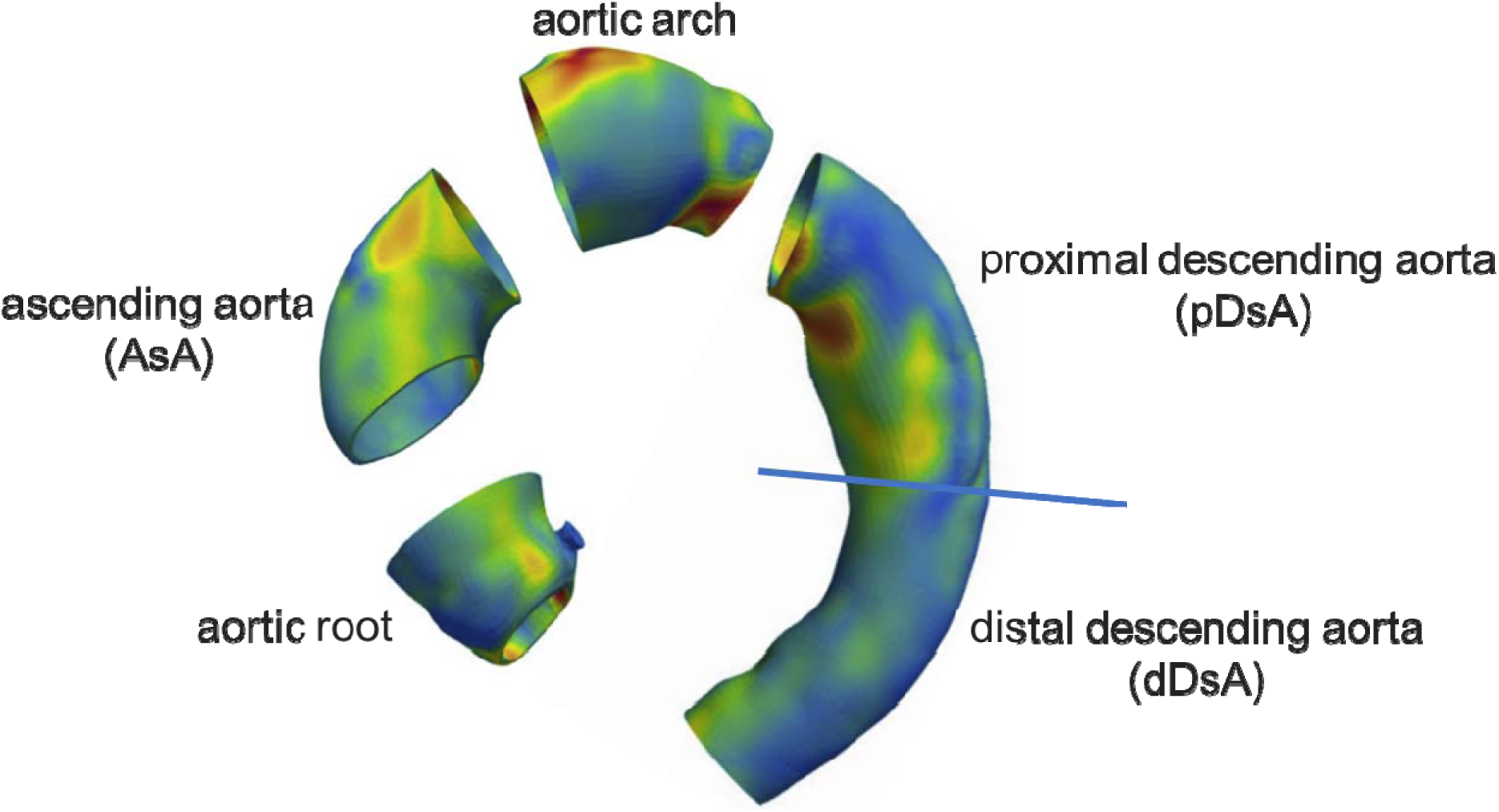
The five regions of the aorta: aortic root, ascending aorta, aortic arch, proximal descending aorta, and distal descending aorta. The aortic root region is cut between the annulus and the sinotubular junction. The ascending aorta is from the sinotubular junction to the brachiocephalic artery. The aortic arch region is cut such that it includes the branches from the brachiocephalic artery to the left subclavian artery. The aorta outlet is cut by a plane approximately passing through the aortic root center. The descending aorta region is cut into a proximal section (from the left subclavian artery to the level of the pulmonary bifurcation), and a distal section (from the level of the pulmonary bifurcation to the termination of the aortic imaging from the CT scan). Stress is color-coded.

**Figure 4:**
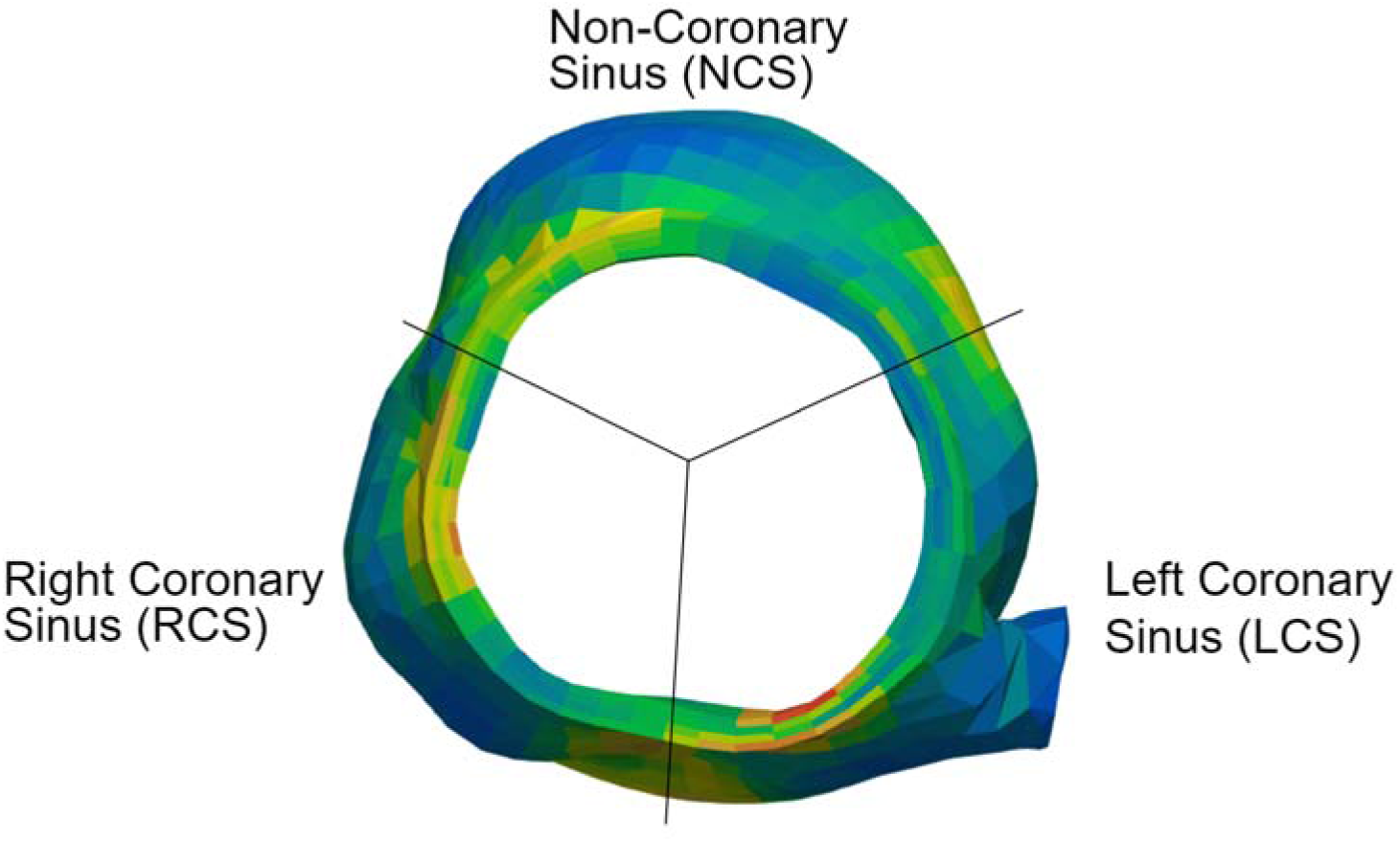
The three segments of the aortic root, viewed from the Sinuses of Valsalva to the sinotubular junction: left coronary sinus, right coronary sinus, and non-coronary sinus. The stress is color-coded.

Paired Student’s t-tests were used to compare the differences in stress between different s ates (preD and postD) and between different regions (in the aorta: Root, AsA, Arch, pDsA, and dDsA, and in the root: LCS, RCS, NCS). For all statistical considerations, a p-value of < 0.05 was deemed to indicate a significant difference. All the data given in the following sections are presented as the mean ± standard deviation.

## 3. RESULTS

### 3.1 Stresses in the aortic wall at preD, postD, and postR states

The stress distributions in the aortic wall of a representative aorta at the preD, postD, and postR states under the normal physiological loading condition of 16 kPa (∼120 mmHg) are shown in Figure 5 from both anterior and posterior views of the aorta. As depicted in Figure 5, the stress color map of the preD state revealed several stress concentration “hotspot” locations in the aortic wall, one of which is present in both the preD state and postD state. The stresses on the graft at the postR state are magnitudes higher than the stresses on other regions because the graft has much smaller thickness compared to the native aortic wall. Since the graft has much higher strength than the native aortic wall tissue, it is not informative to analyze the stress at the postR state.

**Figure 5:**
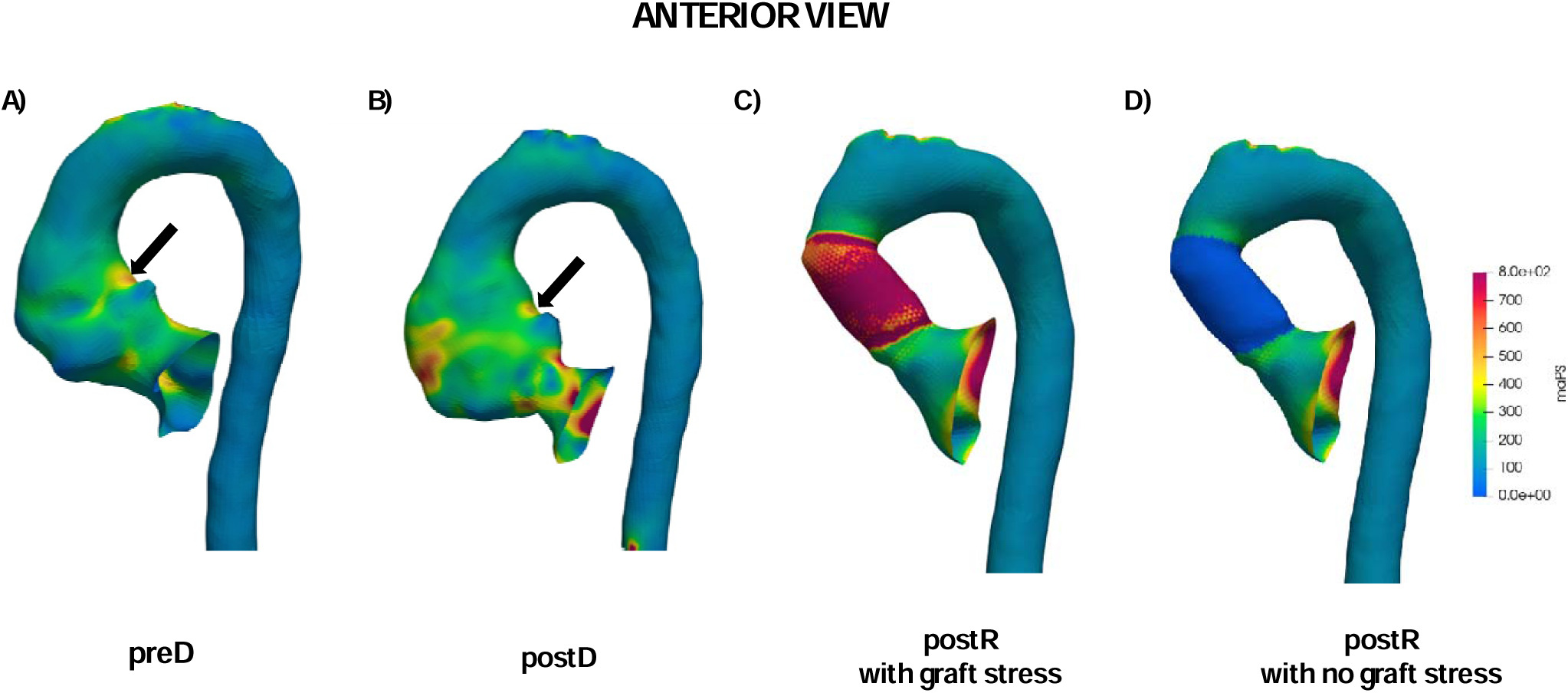
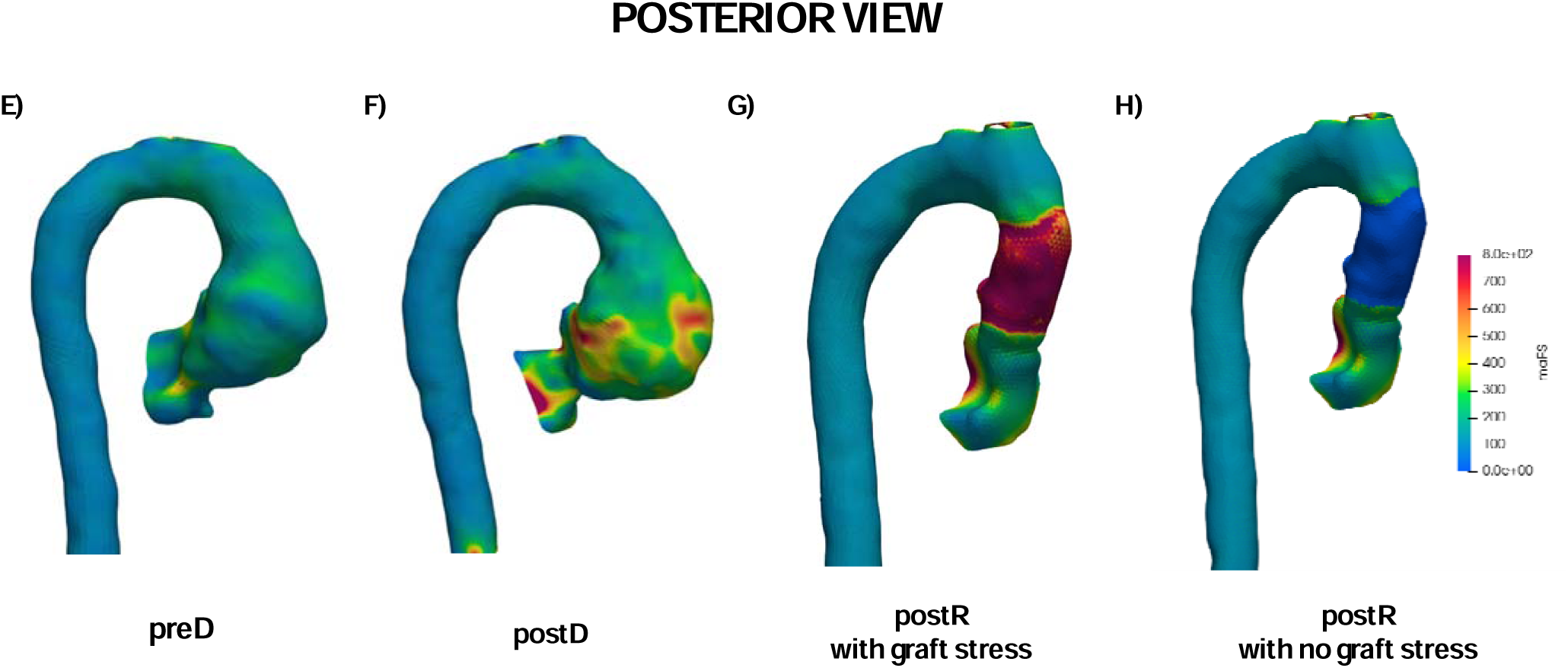
Anterior and posterior views of the aortic wall stress of a representative aorta (Patient #7) at preD, postD and postR states. In D) and H), the stress on the graft is set to 0 to visualize the graft. The stresses on the graft at the postR state are magnitudes higher than the stresses on other regions because the graft has much smaller thickness, compared to the native aortic wall. Since the graft has much higher strength than native aortic wall tissue, it is not informative to analyze the stress at the postR state.

#### 3.1.1 Mean stresses in the aortic wall at preD and postD states

At all regions of the aorta, there was a significant difference (paired t-test, p-value < 0.05) between the preD and postD mean stresses. The mean stresses in the five aorta regions, defined in Figure 3, all increased from the preD state to the postD state, as shown in Figure 6. For example, the mean stress of the aortic root in preD was 161.1±9.2 kPa, which significantly increased to 241.3±21.9 kPa in postD. The mean stresses in the ascending aorta and aortic arch follow the same statistically significant trend. For the descending aorta, the mean stress in the proximal descending aorta was 115.5±4.8 kPa in preD, and significantly increased to 235.8±29.9 kPa in postD, and the same trend was observed in the distal descending aorta segment.

**Figure 6:**
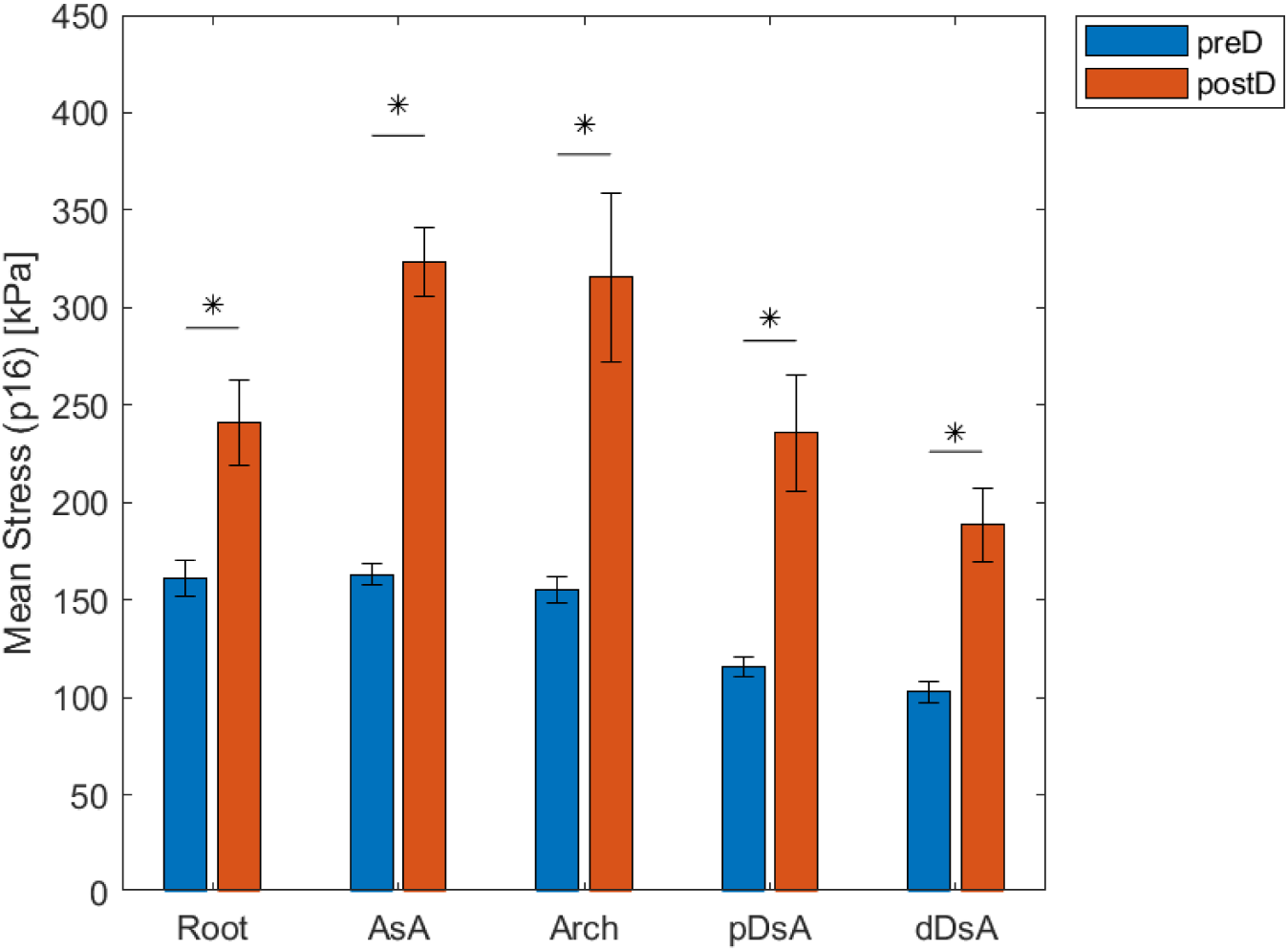
Mean stresses of all patients in the PreD and PostD states, separated by aortic regions. Note: * indicates a statistically significant difference between two groups.

#### 3.1.2 Peak stresses in the aortic wall at preD and postD states

A similar trend occurred in the peak aortic wall stresses in all regions, as shown in Figure 7. In the Root, AsA, pDsA, and dDsA, there was a significant difference (paired t-test, p-value < 0.05) in the peak stresses between the preD and postD states. For example, the peak stresses in the aortic root at the preD state (370.8±35.2 kPa) increased to 681.9±100.2 kPa in the postD state. The peak stress in the pDsA region was 230.6±13.8 kPa at the preD state, which significantly increased to 714.3±144.69 kPa in the postD state.

**Figure 7:**
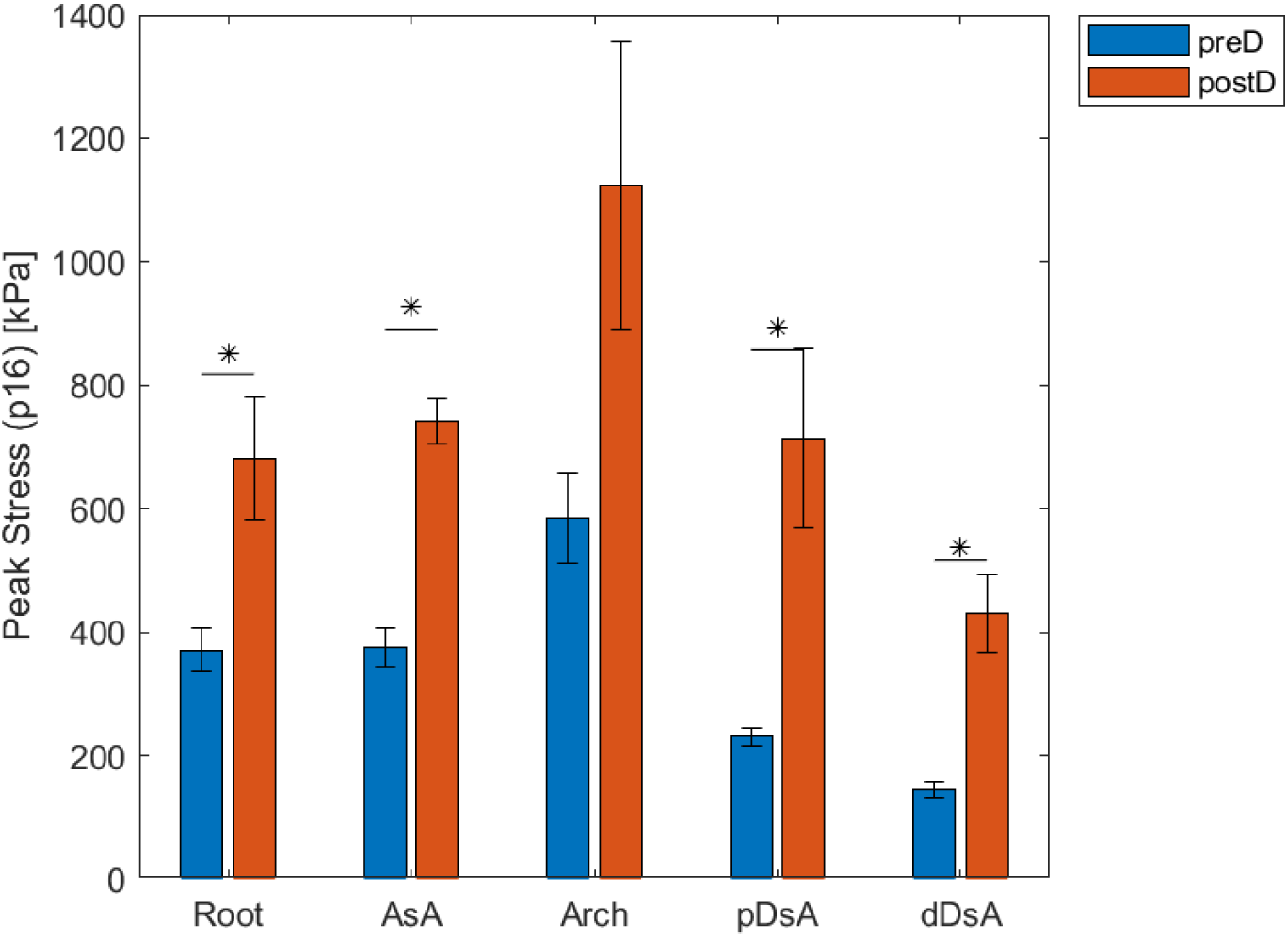
Peak stresses of all patients in the PreD and PostD states, grouped by aortic regions.

For the stress changes in the descending aorta, it is worth noting that only three patients had a dissection that was limited to the ascending aorta. In one such patient, the peak stress in the pDsA was not changed substantially (i.e., 236.17 kPa in the preD state, and 344.08 kPa in the postD state), and similarly, the peak stress in the dDsA did not change substantially (i.e., 119.87 kPa in the preD state, and 132.41 kPa in the postD state). The mean stresses in the pDsA and dDsA followed a similar trend (i.e., for pDsA, 113.76 kPa in the preD state and 114.80 kPa in the postD state; for dDsA, 96.36 kPa in the preD state, and 102.60 kPa in the postD state). This may indicate that the increased stress in the descending aorta after dissection in this dataset is largely due to the dissection propagating to the descending aorta.

Interestingly, there was no significant difference between peak stresses in the preD and postD states for the Arch. When comparing between the regions of the aorta at the preD state, we found that there was a significant difference in the peak stresses for all comparisons except one: There was no significant difference between the peak stresses of the Root and the AsA in the preD state.

Overall, the ascending aorta (consisting of three regions: Root, AsA, and Arch, see Figure 3) experienced higher mean and peak stresses than the descending aorta (pDsA and dDsA) in the preD state, mostly due to the formation of aneurysm at the ascending aorta. For example, in the preD state, the mean stress was 161.1 kPa in the Root, 162.9 kPa in the AsA region, and 155.2 kPa in the Arch, whereas the mean stress was 115.2 kPa in the pDsA region, and 102.7 kPa in the dDsA region. Again, in the preD state, the peak stress was 370.8 kPa in the Root, 374.6 kPa in the AsA region, and 585.5 kPa in the Arch, whereas the peak stress is 230.6 kPa in the pDsA region and 145.4 kPa in the dDsA region.

### 3.2. Stresses of the LCS, RCS, and NCS sections of the aortic root at preD and postD states

The mean stress analysis results of the three aortic root sections (LCS, RCS, and NCS) can be seen in Figure 8. There were no significant differences between the mean stresses of the three regions in the preD state. There was again a significant increase in mean stresses from the preD state to the postD state, and the mean stresses decreased after surgical repair. In the preD state, the LCS, RCS and NCS had mean stresses of 173.4±13.9 kPa, 168.6±11.1 kPa, and 149.0±10.8 kPa, respectively. The NCS experienced the highest increase in stress from the preD state to the postD state compared to the LCS and RCS, increasing an average of 111.6 kPa whereas the LCS increased 55.6 kPa, and the RCS increased 70.7 kPa.

**Figure 8:**
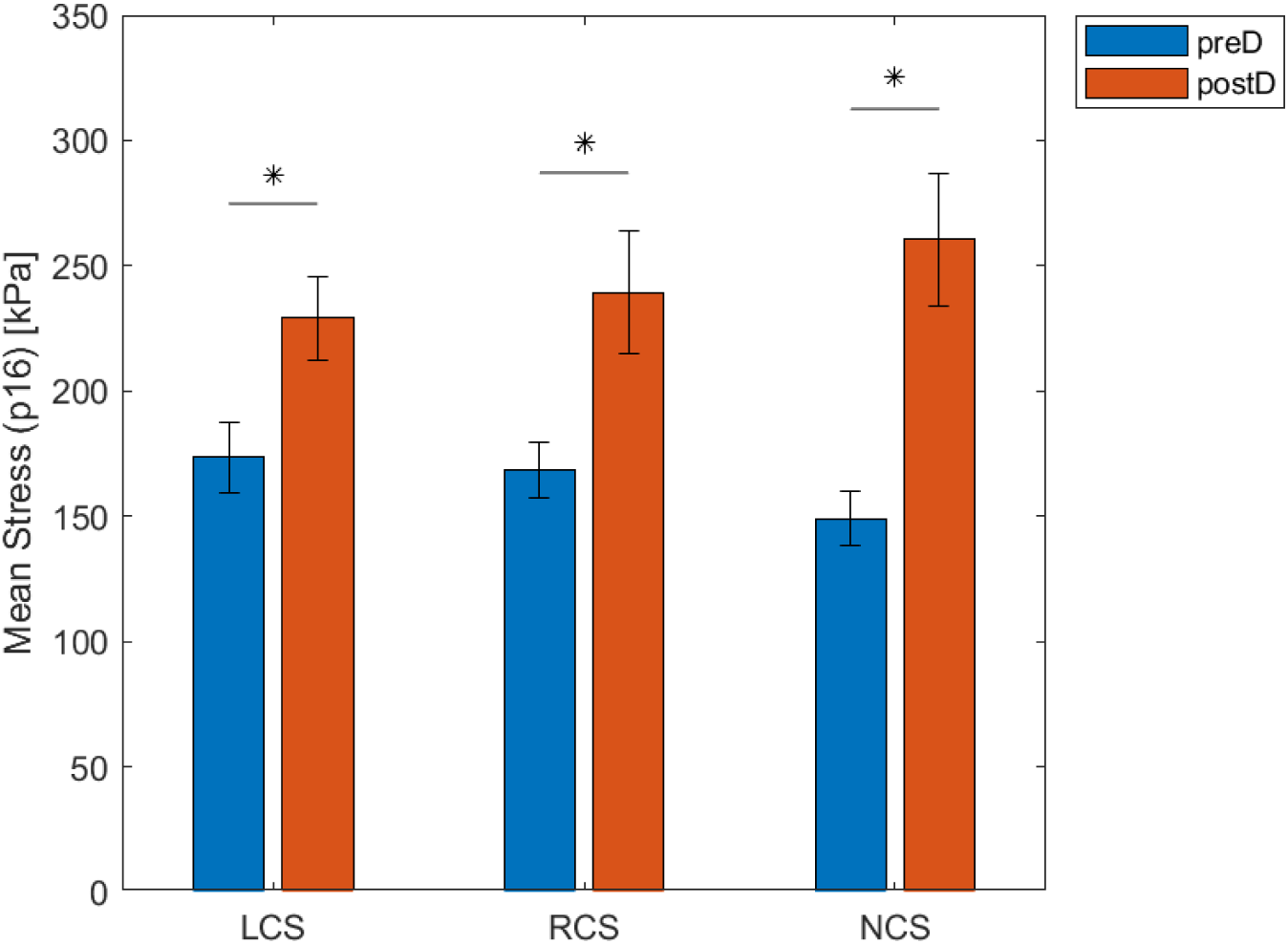
Mean stresses of all patients in the PreD and PostD states, separated by aortic root regions.

The peak stress analysis results of the three aortic root sections (LCS, RCS, and NCS) can be seen in Figure 9. In the preD state, there is no significant difference in the peak stresses between the LCS and RCS and between the LCS and NCS, but there is a significant difference in the peak stresses between the RCS and NCS. There was an increase in peak stress from the preD state to the postD state. For example, the LCS had a peak stress of 355.8 kPa±40.2 kPa in the preD state, and the peak stress increased to 562.5±51.4 kPa in the postD state. There was also a significant difference in the peak stresses between the preD and postD states in the LCS and NCS.

**Figure 9:**
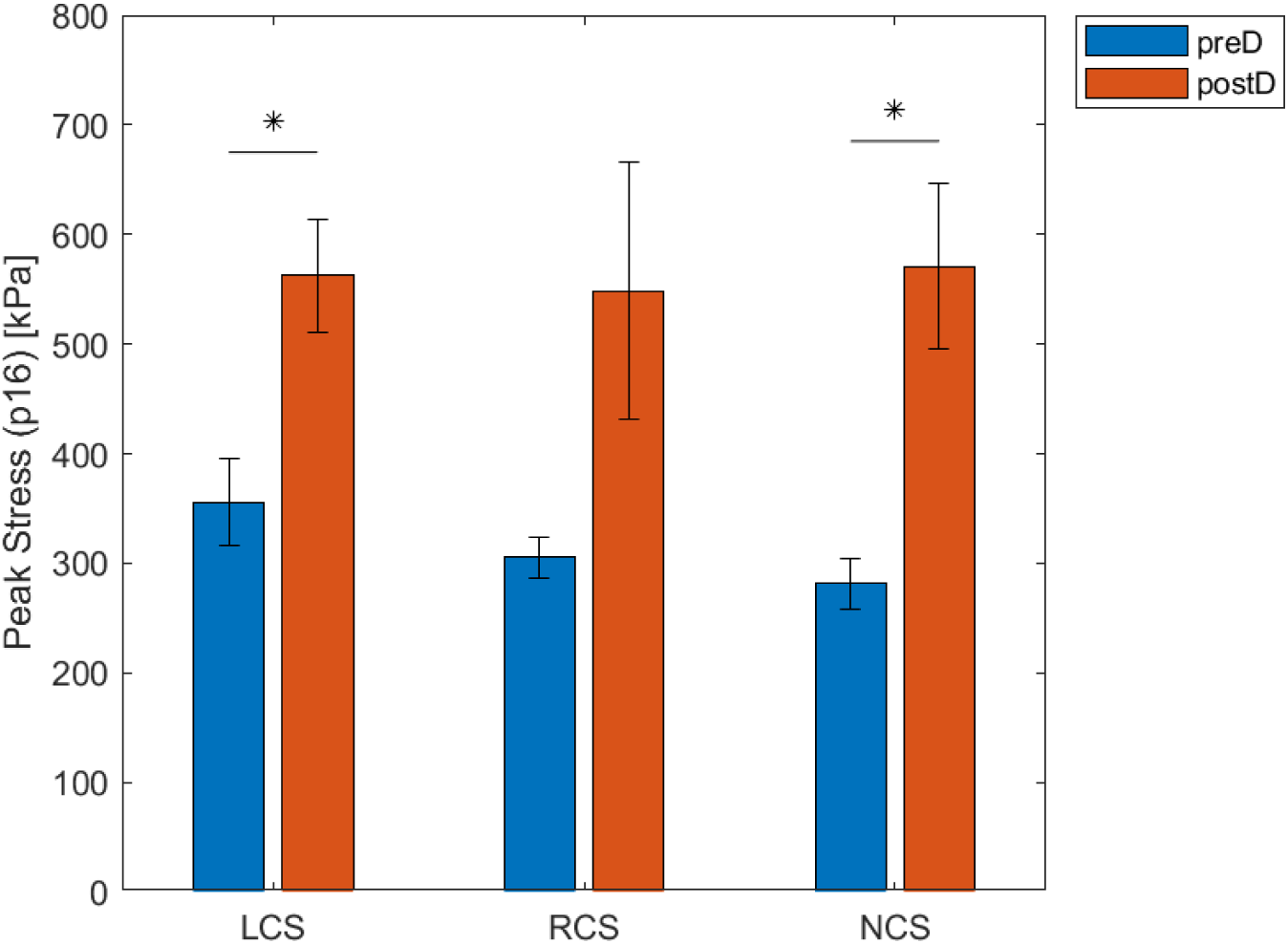
Peak stresses of all patients in the PreD and PostD states, grouped by aortic root regions.

### 3.3. Stresses of the aorta under hypertensive blood pressures

The blood pressure of a patient could temporarily increase as a response to emotional stress, anxiety, or carrying heavy objects. For example, weightlifting could cause extreme (>200 mmHg or 26kPa) elevations in blood pressure leading to aortic rupture [35]. We further performed stress analysis at different blood pressure levels, i.e., investigating how the stresses of different aorta regions would change at different blood pressure levels: 16 kPa (normotensive), 18 kPa (hypertension stage-1), and 20 kPa (hypertension stage-2). The results are reported in Tables 2 and 3. The stresses at the hypertensive blood pressures in this analysis are the lower bounds of the *in vivo* stresses for two reasons: (1) the stresses were computed based on the SDA principle (Section 2.3) with geometries reconstructed from CT images at a normotensive pressure, and (2) the actual aorta diameter of a patient may increase under a pressure higher than the pressure captured during CT imaging, resulting in the actual stresses being higher than the SDA-computed stresses. The significantly elevated stresses (lower-bounds) at higher blood pressure levels (Tables 2 and 3) indicate higher risk of rupture if patients were left untreated.

**Table 2:** the mean stresses (kPa) of different regions at three blood pressure levels.

|  | Pressure = 16 kPa |  | Pressure = 18 kPa |  | Pressure = 20 kPa |  |
| --- | --- | --- | --- | --- | --- | --- |
|  | preD | postD | preD | postD | preD | postD |
| AsA | 162.9 ± 5.6 | 323.6 ± 17.8 | 183.3 ± 6.4 | 364.7 ± 20.4 | 203.5 ± 6.6 | 404.5 ± 22.1 |
| Arch | 155.2 ± 6.8 | 315.5 ± 43.1 | 173.6 ± 7.8 | 343.4 ± 53.7 | 192.5 ± 8.3 | 379.7 ± 58.5 |
| pDsA | 115.2 ± 4.8 | 235.8 ± 29.9 | 131.2 ± 5.7 | 265.5 ± 34.3 | 146.2 ± 5.8 | 292.9 ± 37.7 |
| dDsA | 102.7 ± 5.2 | 188.5 ± 19.3 | 115.9 ± 5.8 | 216.5 ± 22.9 | 128.6 ± 6.4 | 236.2 ± 24.1 |
| Root | 161.1 ± 9.2 | 241.3 ± 21.9 | 181.5 ± 10.5 | 265.5 ± 22.0 | 198.6 ± 12.0 | 296.5 ± 24.4 |
| LCS | 173.4 ± 13.9 | 229.0 ± 16.9 | 196.0 ± 14.3 | 257.6 ± 20.5 | 241.2 ± 18.5 | 285.2 ± 22.3 |
| RCS | 168.6 ± 11.1 | 239.3 ± 24.4 | 189.3 ± 12.5 | 261.8 ± 20.7 | 206.3 ± 14.5 | 289.8 ± 20.0 |
| NCS | 149.0 ± 10.8 | 260.5 ± 26.4 | 165.6 ± 10.6 | 287.7 ± 27.2 | 182.2 ± 12.0 | 325.8 ± 32.6 |

**Table 3:**
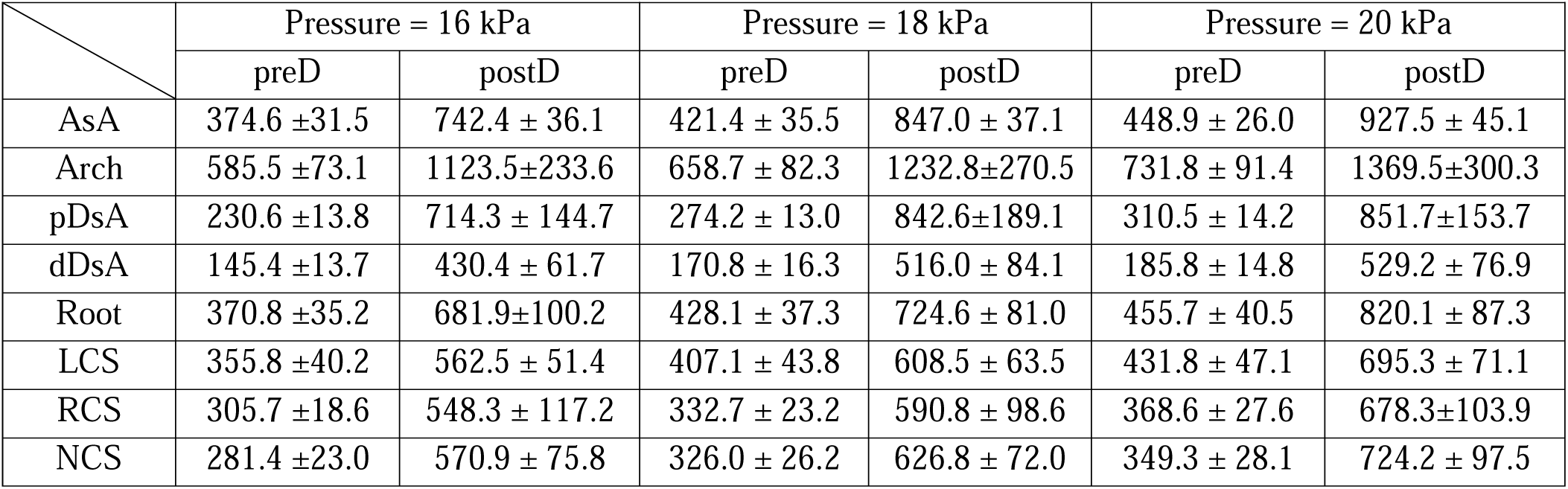
the peak stresses (kPa) of different regions at three blood pressure levels.

|  | Pressure = 16 kPa |  | Pressure = 18 kPa |  | Pressure = 20 kPa |  |
| --- | --- | --- | --- | --- | --- | --- |
|  | preD | postD | preD | postD | preD | postD |
| AsA | 374.6 ±31.5 | 742.4 ± 36.1 | 421.4 ± 35.5 | 847.0 ± 37.1 | 448.9 ± 26.0 | 927.5 ± 45.1 |
| Arch | 585.5 ±73.1 | 1123.5±233.6 | 658.7 ± 82.3 | 1232.8±270.5 | 731.8 ± 91.4 | 1369.5±300.3 |
| pDsA | 230.6 ±13.8 | 714.3 ± 144.7 | 274.2 ± 13.0 | 842.6±189.1 | 310.5 ± 14.2 | 851.7±153.7 |
| dDsA | 145.4 ±13.7 | 430.4 ± 61.7 | 170.8 ± 16.3 | 516.0 ± 84.1 | 185.8 ± 14.8 | 529.2 ± 76.9 |
| Root | 370.8 ±35.2 | 681.9±100.2 | 428.1 ± 37.3 | 724.6 ± 81.0 | 455.7 ± 40.5 | 820.1 ± 87.3 |
| LCS | 355.8 ±40.2 | 562.5 ± 51.4 | 407.1 ± 43.8 | 608.5 ± 63.5 | 431.8 ± 47.1 | 695.3 ± 71.1 |
| RCS | 305.7 ±18.6 | 548.3 ± 117.2 | 332.7 ± 23.2 | 590.8 ± 98.6 | 368.6 ± 27.6 | 678.3±103.9 |
| NCS | 281.4 ±23.0 | 570.9 ± 75.8 | 326.0 ± 26.2 | 626.8 ± 72.0 | 349.3 ± 28.1 | 724.2 ± 97.5 |

A representative aorta at the postD state is illustrated in Figure 10 in which Figure 10A&E show anterior and posterior views of true and false lumens, and the remaining 3 columns show the stresses on the aortic wall under pressure levels of 16 kPa (∼120 mmHg), 18 kPa (∼135 mmHg), and 20 kPa (∼150 mmHg), respectively.

**Figure 10:**
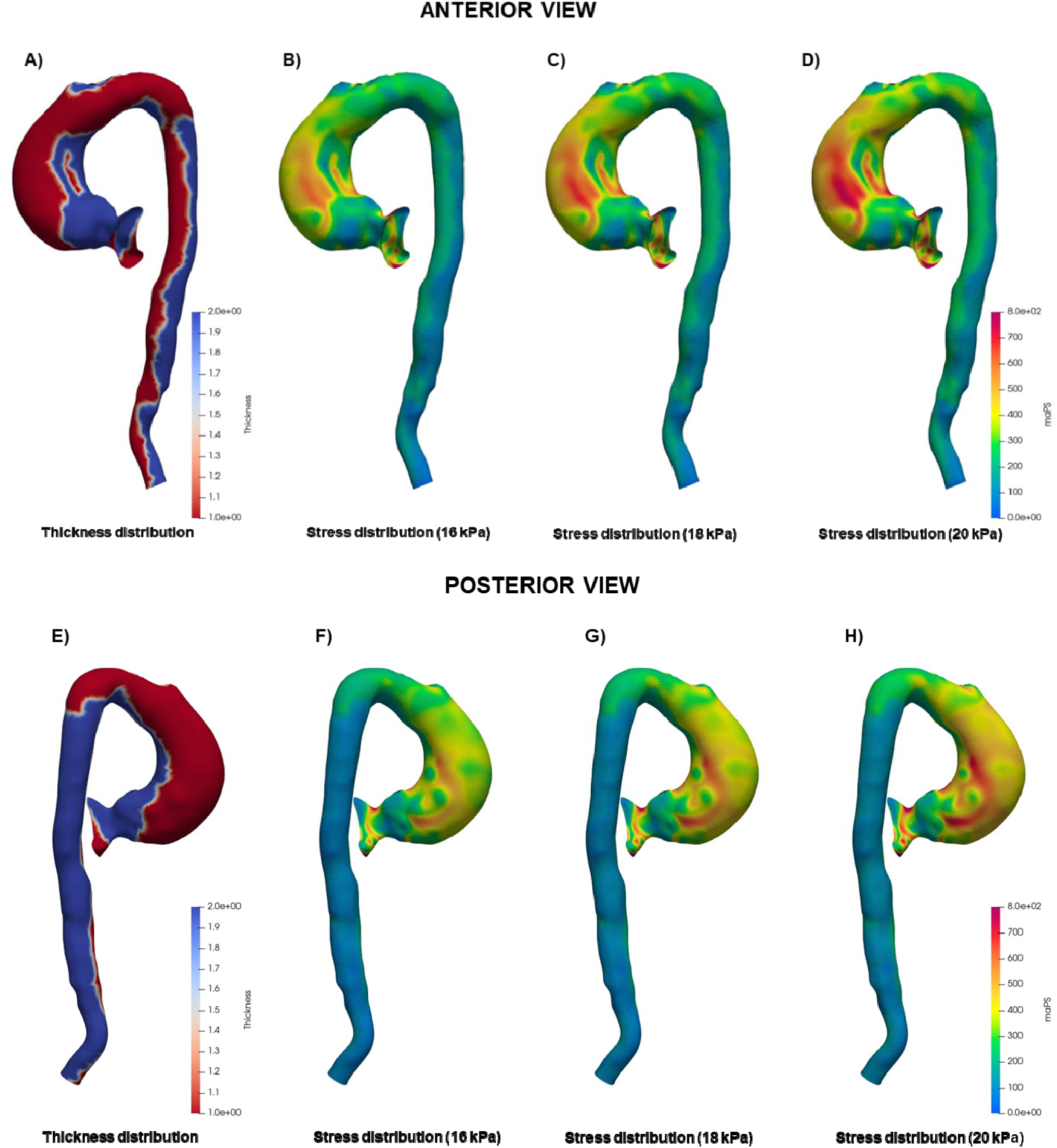
Anterior and posterior views of the stress distribution of a representative aorta at the postD state at 16, 18, and 20 kPa. A&E show anterior and posterior views of true and false lumens (color represents the thickness), B&F show the stresses distribution under 16 kPa (∼120 mmHg), C&G show the stresses distribution under 18 kPa (∼135 mmHg), D&H show the stresses distribution under 20 kPa (∼150 mmHg).

### 3.4. Geometry-Stress analysis of the aorta at the preD, postD, and postR states

We investigated the correlation between stress and two commonly used geometric features (curvature and radius). We recall that, for each patient, the stress analysis was performed using a solid element mesh that was obtained by offsetting a surface mesh with a prescribed thickness; at each element of the solid mesh, the maximum of the absolute values of the principle stresses was obtained, and this value is simply called “stress” in this study; each element of the surface mesh is associated with a stress value that is the stress averaged across the thickness. Because geometric features are usually defined on nodes, for each node of the surface mesh, we obtained a stress value by linearly interpolating the stresses of the neighbor elements. Thus, for each patient, we obtained node-stress pairs on a surface mesh, and at each node, we obtained mean curvature and radius as geometric features associated with the node. Because the shape of the aorta is not a cylindrical tube, there exist at least three different definitions of radius, and each of the radius definitions needs the centerline of the aorta. The first definition (def. 1) is the Maximum Inscribed Sphere Radius, which can be determined through the following steps: start by creating a small sphere centered at a point on the centerline, gradually increase the radius of the sphere as much as possible while ensuring it remains within the aorta, and finally, measure the radius of the largest sphere that fits entirely inside the aorta. The second definition (def. 2) is Mean Radius of Cross Section, which can be determined through the following steps: get the cross section passing a point on the centerline, measure the area of the cross section, and compute the radius by the equation: area = 3.14×radius×radius. The third definition (def. 3) is Max Radius of Cross Section, which is the max radius of a cross section. For each node on the aortic wall, the nearest point on the centerline can be found, and then the radius associated with the node can be calculated using def. 1, def. 2, or def. 3.

In this study, centerlines and geometric features were obtained by using the VMTK library [36]. We calculated Pearson correlation coefficients between stress and each of the geometric features when blood pressure is 16 kPa. The results are reported in Table 4.

**Table 4:** The correlations between stress and each of the geometric features.

| Patient ID | State | Curvature | Radius (def. 1) | Radius (def. 2) | Radius (def. 3) |
| --- | --- | --- | --- | --- | --- |
| 1 | preD | 0.2663 | 0.3168 | 0.3669 | 0.3748 |
|  | postD | 0.1073 | 0.1392 | 0.0677 | 0.2193 |
|  | postR | 0.0531 | -0.0913 | -0.1009 | -0.1194 |
| 2 | preD | 0.4160 | 0.6106 | 0.7328 | 0.7412 |
|  | postD | 0.3061 | 0.7095 | 0.7263 | 0.7566 |
|  | postR | 0.0652 | 0.4408 | 0.5335 | 0.4516 |
| 3 | preD | 0.2350 | 0.1041 | 0.3761 | 0.3398 |
|  | postD | 0.0822 | 0.4673 | 0.1979 | 0.4715 |
| 4 | preD | 0.1964 | 0.3038 | 0.5095 | 0.4919 |
|  | postD | 0.0757 | 0.2187 | -0.0336 | 0.1992 |
| 5 | preD | 0.2982 | 0.2710 | 0.4514 | 0.4667 |
|  | postD | 0.0036 | 0.2005 | 0.0768 | 0.1137 |
| 6 | preD | 0.2971 | 0.4053 | 0.5769 | 0.6118 |
|  | postD | -0.0481 | 0.2259 | 0.0943 | 0.1065 |
| 7 | preD | 0.2864 | 0.4081 | 0.4999 | 0.5002 |
|  | postD | 0.2640 | 0.5339 | 0.5343 | 0.5431 |
|  | postR | 0.0500 | 0.1829 | 0.3088 | 0.2675 |
| 8 | preD | 0.2687 | 0.5018 | 0.6041 | 0.6162 |
|  | postD | 0.1590 | 0.4295 | 0.5023 | 0.5104 |
|  | postR | 0.0402 | 0.2765 | 0.3643 | 0.3285 |
| (average) | preD | $0.2830 \pm 0.0596$ | $0.3652 \pm 0.1438$ | $0.5147 \pm 0.1144$ | $0.5178 \pm 0.1246$ |
| | postD | $0.1187 \pm 0.1131$ | $0.3656 \pm 0.1873$ | $0.2708 \pm 0.2594$ | $0.3650 \pm 0.2226$ |
| | postR | $0.0521 \pm 0.0089$ | $0.2022 \pm 0.1930$ | $0.2764 \pm 0.2330$ | $0.2320 \pm 0.2135$ |

The correlation between stress and radius can be explained by the Laplace law, which states that the hoop stress is equal to pressure × radius / thickness for a thin-walled cylindrical pressure vessel. At the postD and postR states, the thickness of the aortic wall is smaller in the regions of dissection and the region of graft, which may affect the correlation analysis. Therefore, we calculated the “adjusted” stress defined by stress × (thickness/2), where 2 (mm) is the normal wall thickness, and we subsequently computed the correlation between the adjusted stress and each of the geometric features at the postD and postR states. The results are reported in Table 5, which do not show any large improvement compared to the results in Table 4.

**Table 5:**
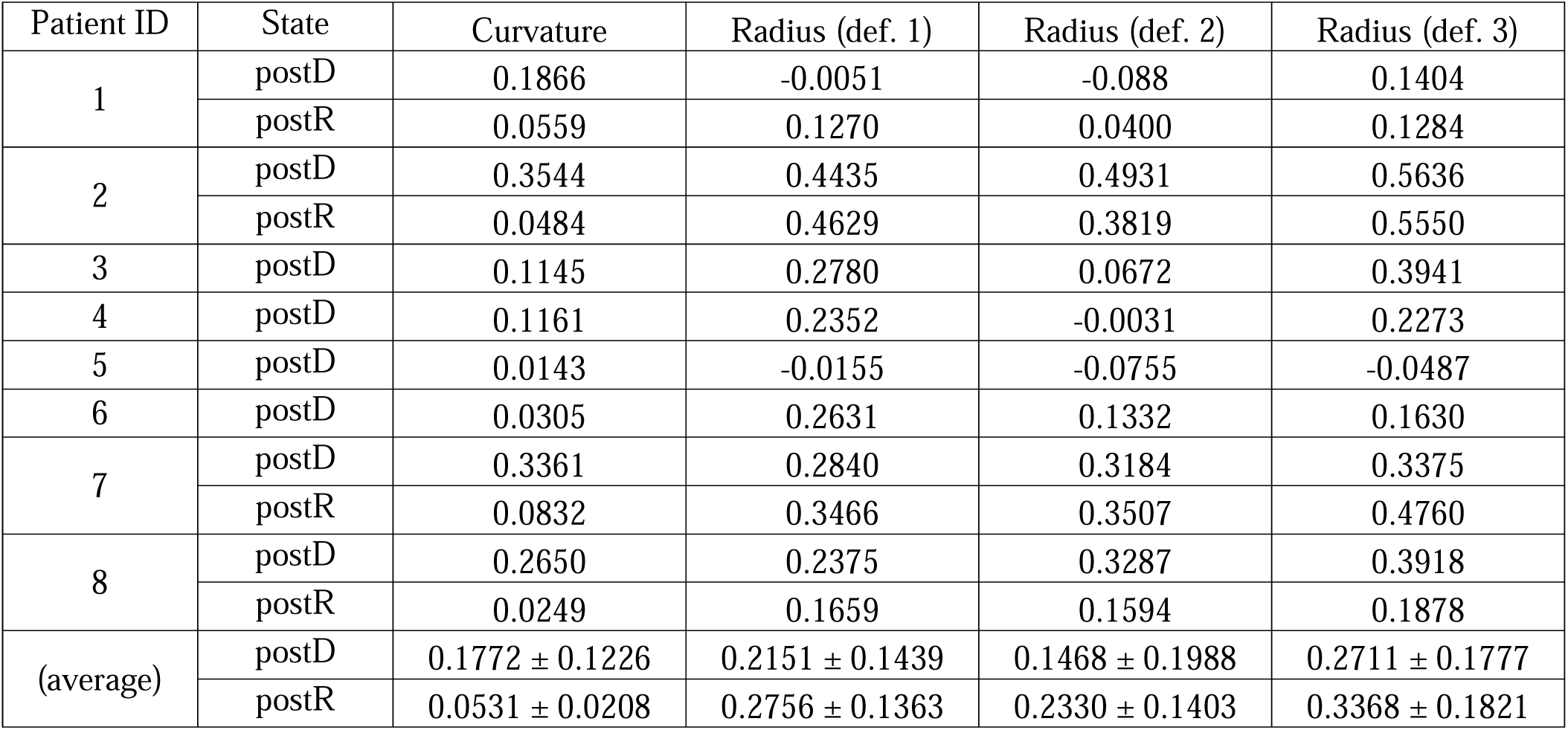
The correlations between the adjusted stress and each of the geometric features.

From Table 2 and Table 3, it can be observed that the stresses at both the ascending and descending aorta increased from the preD state to the postD state on average. To explain these increases, we conducted aorta diameter analysis of the eight patients. For ascending aorta, we calculated the change in maximum radius from preD to postD states, i.e., max radius at postD – max radius at preD, and the results are in Table 6. All eight patients had ascending aorta dissection. For descending aorta, we calculated the change in average radius from preD to postD states, i.e., average radius at postD – average radius at preD, and the results are in Table 7. Five Patients (1, 2, 4, 6, and 8) had both ascending and descending aorta dissection, and the other three patients (3, 5, and 7) had dissection at the ascending aorta but did not have dissection at the descending aorta.

**Table 6:** The change in ascending aorta radius (mm) from preD to postD (negative means decrease)

| Patient ID | Ascending aorta dissection | Change in radius (def. 1) | Change in radius (def. 2) | Change in radius (def. 3) |
| --- | --- | --- | --- | --- |
| 1 | Yes | 0.9003 | 1.4378 | 1.3418 |
| 2 | Yes | 4.1745 | 5.2963 | 4.3476 |
| 3 | Yes | 2.0533 | 2.2718 | -0.2008 |
| 4 | Yes | 2.8366 | 2.8738 | 1.3868 |
| 5 | Yes | 0.719 | 2.9787 | 1.717 |
| 6 | Yes | 6.0884 | 5.762 | 4.8255 |
| 7 | Yes | 4.4837 | 5.1707 | 5.761 |
| 8 | Yes | 3.9506 | 5.3158 | 6.1992 |

**Table 7:**
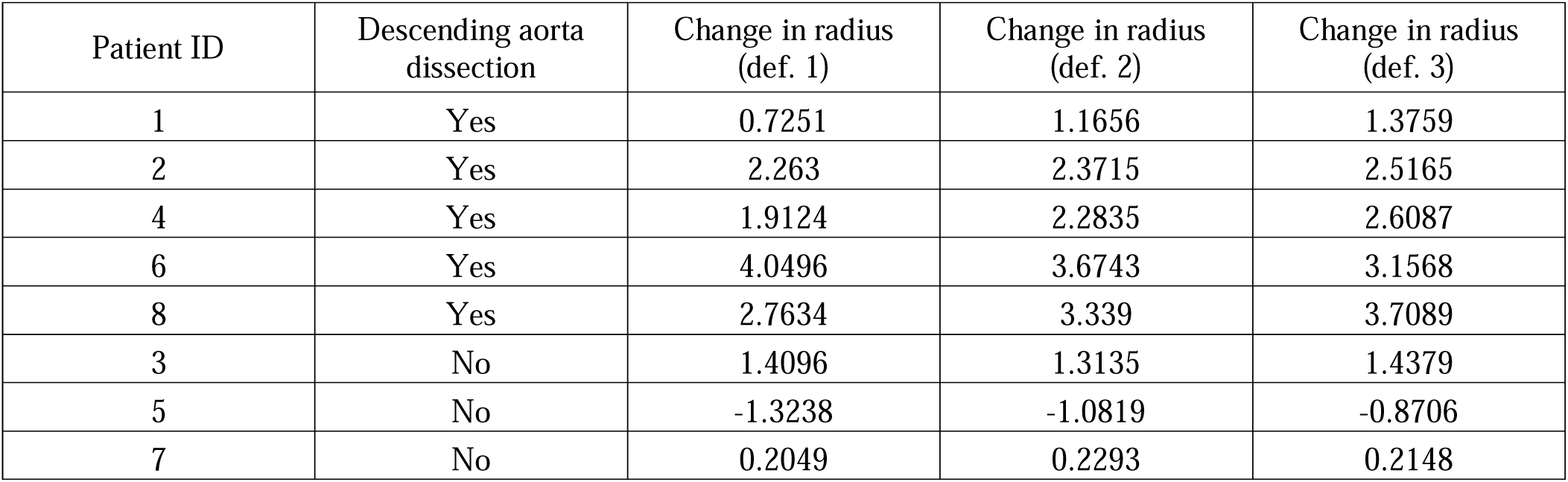
The change in descending aorta radius (mm) from preD to postD (negative means decrease)

| Patient ID | Descending aorta dissection | Change in radius (def. 1) | Change in radius (def. 2) | Change in radius (def. 3) |
| --- | --- | --- | --- | --- |
| 1 | Yes | 0.7251 | 1.1656 | 1.3759 |
| 2 | Yes | 2.263 | 2.3715 | 2.5165 |
| 4 | Yes | 1.9124 | 2.2835 | 2.6087 |
| 6 | Yes | 4.0496 | 3.6743 | 3.1568 |
| 8 | Yes | 2.7634 | 3.339 | 3.7089 |
| 3 | No | 1.4096 | 1.3135 | 1.4379 |
| 5 | No | -1.3238 | -1.0819 | -0.8706 |
| 7 | No | 0.2049 | 0.2293 | 0.2148 |

### 3.5. Aorta diameter analysis for the patients #1, 2, 7, 8 at the preD, postD, and postR states

For patients # 1, 2, 7, 8 (see Table 1), the postR CT image data were available. For each of these patients at the postR state, part of the ascending aorta was replaced by a graft, and the stress on the graft was the highest among different regions because graft thickness is around 0.3mm [24], much smaller than the aortic wall (2mm for norma wall, and 1mm for dissected wall). Compared to the native aorta tissues, the graft has much higher strength and will not rupture under hypertensive blood pressures; therefore, stress analysis is not informative for these patients. Instead, we conducted aorta diameter analysis as shown in Figure 11.

**Figure 11.**
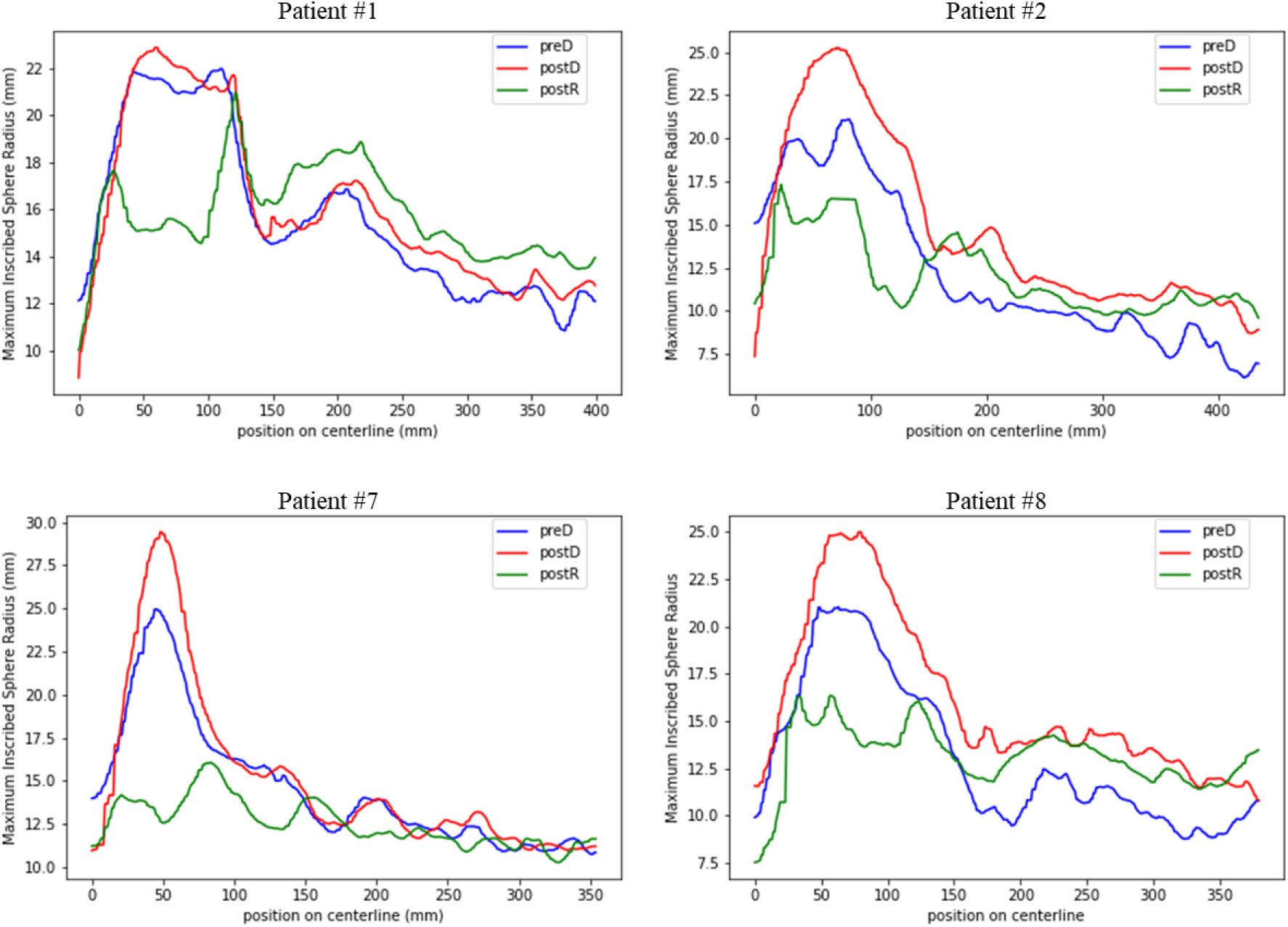
Aorta diameter analysis for the four patients (1,2,7,8). Each curve shows the change of radius (def. 1) along the centerline. Position 0 on the centerline is at the center of aortic annulus. The ascending aortic aneurysm has the largest radius on each curve.

**Figure 12:**
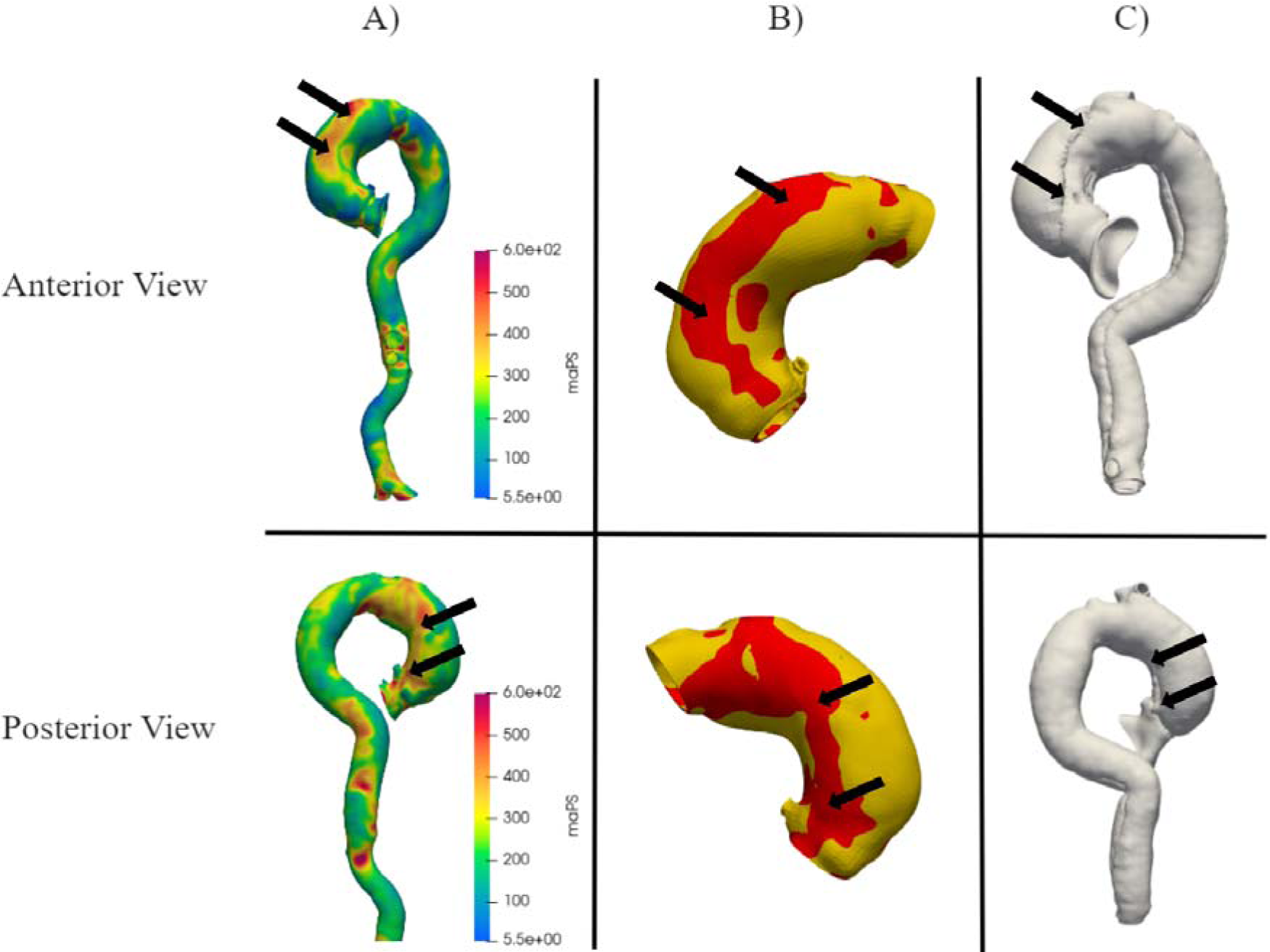
Anterior and posterior views of a representative aorta from Patient #1 with (A) the PreD stress distribution, (B) a threshold image of just the ascending aorta, where the red region represents the stresses greater than the 75th percentile, and (C) the PostD state. The arrows show areas of stress concentration and the dissection borders. For a better visualization, two separate meshes are used for the true and false lumens in (C).

## 4. DISCUSSION

### 4.1. High aortic wall stress at the postD state (necessity of surgery)

Using 16 kPa (∼120mmHg) as the normal systolic blood pressure, the stresses at the postD state are significantly higher than those at the preD state. For example, the mean stress in the AsA region was 162.9±5.6 kPa at the preD state and 323.6±17.8 kPa at the postD state under 16 kPa systolic blood pressure. The increase in stress leads to an increased risk of rupture/dissection. To quantify the risk of rupture, we calculated the probability index of rupture, which was developed in our previous work [37], and the aneurysm rupture probability of each of the eight patients at the postD state under the 16 kPa systolic blood pressure is higher than 0.9, indicating imminent rupture if left untreated.

### 4.2. Surgical repair restores aorta diameter close to normal (effectiveness of surgery)

The ascending aorta aneurysms of the eight patients were replace by grafts. From Figure 11, it can be seen that surgeries reduced the diameters of the ascending aorta, although the dilated aortic arch of Patient #1 was not fully replaced by a graft. However, the descending aorta diameters of three patients (1, 2, and 8) at the postR and postD states are larger than those at the preD stage, and this is because the three patients had dissections at both the ascending aorta and the descending aorta. Right after dissection happens at the descending aorta, the outer wall has smaller thickness and therefore has higher stress, leading to wall expansion until force equilibrium is reached. At the descending aorta, the stress at postD (and postR) is higher than stress at preD because of the increase in diameter and decrease in thickness. Although the descending aorta stress is elevated, the diameter of the descending aorta is still less than 40 mm, which is considered low risk and therefore not treated [38].

### 4.3 Positive but weak correlations between stress and diameter/curvature

From Table 4 and Table 5, it can be seen that there are positive correlations between stress and each of the geometric features (i.e., curvature and radius) in general. At the preD state, the correlation between stress and Max Radius of Cross Section is relatively high (r=0.5178 ± 0.1246), and the correlation between stress and curvature is weak (r=0.2830 ± 0.0596). At the postD and postR states, the correlations are weak (average r << 0.5). The results of this study are consistent with the results of our previous study of aneurysm rupture risk and aorta geometry [39]. In the previous study, we used a synthetic dataset of aortic aneurysm geometries (no dissection) generated by a statistical shape model, and performed rupture risk analysis; the results show that rupture risk is only weakly correlated with diameter and curvature, and it is determined by the whole geometry of the aorta when other conditions are fixed.

The results of the correlation analysis indicate that stress is not fully determined by radius and curvature at the preD, postD, and postR states. Therefore, FE stress analysis is necessary to obtain the full stress distribution, from which the probabilistic rupture risk can be calculated [37].

### 4.4 Comparing between regions of the aorta

The stresses in each of the ascending aorta regions (including the Root, AsA, and Arch) were consistently higher than those in the descending aorta (including the pDsA and the dDsA). At the preD state with 16 kPa pressure, there was no significant difference in mean stress between the regions of the ascending aorta (including Root, AsA and Arch). When comparing the peak stresses, only the Root and the AsA had no significant difference.

We observed that stresses at Arch are relatively high. While there was no significant difference in mean stress between the Root and the AsA, the peak stresses in the Arch were significantly higher than those in both the root and the AsA. Since curvature and diameter are only weakly correlated with stress, the relatively high stress at Arch can only be explained by its relatively complex shape, compared to a simple tube, which also highlights the necessity of FE stress analysis.

The increase in stresses in the ascending aorta and the descending aorta from the preD state to the postD state could be partially explained by the increase in diameter and the decrease in thickness. From Table 6, it can be observed that the maximum diameter of the ascending aorta increased for every patient from the preD state to the postD state. For each patient, right after ascending aorta dissection, the diameter of the ascending aorta would increase in order to reach force equilibrium. The stresses at the ascending aorta would increase as the result of increase in diameter and decrease in thickness, and similar explanation can be applied to the descending aorta, according to Table 7.

### 4.5 Possible correlation between dissection and high-stress spots

From the FE stress analysis, at the 16 kPa (∼120mmHg) physiological loading condition, the mean stress in the PreD state is 161.1±9.2 kPa in the aortic root, 162.9±5.6 kPa in the ascending aorta, and 155.2±6.8 kPa in the aortic arch. The mean stresses in these three regions were significantly higher (p < 0.001) compared to the descending aorta, which had a mean stress of 111.2±4.8 kPa in the proximal descending aorta and 102.7±5.2 kPa in the distal descending aorta. Furthermore, we identified stress concentration regions by defining them as areas with stress values greater than the 75th percentile in the aortic wall. Interestingly, for one of the patients, these high-stress regions exhibited strong correlations with the actual dissection borders observed in patients at the post-dissection stage (Figure 2).

### 4.6 Limitation and future work

This study has several limitations, and we intend to resolve the limitations in our future work. Firstly, the wall thickness was assumed to be uniform. The thickness of the aortic wall without dissection was assumed to be 2 mm [40, 41], while the thickness of the dissection wall was set to be half of the normal value according to clinical observations from Dr. Elefteriades. Prior reports have shown that the aortic arch has a wide variation in wall thickness, and there is a thicker wall in the sharp curves between aortic branches [42]. This thicker wall of aorta arch would act as a natural reinforcement to reduce the high peak stresses we observed in the simulation, and it is likely that the true stresses in the aortic arch are lower than the calculated stresses by FEA. More accurate wall thickness can be obtained by using both the contrast and non-contrast CT images, if both are available: the inner wall will be revealed by the contrast CT image, and the outer wall will be revealed by the non-contrast CT image. Secondly, the normal systolic pressure was assumed to be a fixed number, instead of a range of feasible pressure values. An interesting future study could be the investigation of rupture/dissection probability with patient-specific blood pressures, which could provide useful information for blood pressure control to avoid rupture while waiting for surgery. This study did not investigate dissection progression. To investigate dissection progression after a dissection tear occurs, the fluid-structure interaction (FSI) approach could be used. However, it would be difficult to obtain image data at the exact dissection tear formation state. This study did not consider the effect of heart and aorta motion [43] because we only obtained CT data at one single cardiac phase, and this limitation could be resolved by obtaining multi-phase data. In addition, our future work will include machine learning-based FEA [44, 45] to facilitate a fast and accurate estimate of aortic wall stress and allow for patient-specific diagnosis and prediction of aortic dissection/rupture which could be used in the office in clinical practice.

## 5. CONCLUSIONS

We performed patient-specific stress and geometry analysis of aortic dissection at three time points: preD, postD, and postR. By analyzing mean and peak stress values as well as geometric features across different regions and states, we observe distinct patterns that highlight the mechanical response of the aorta to dissection and subsequent repair.

## ACKNOWLEDGEMENTS

This study was supported in part by the NIH grant R01HL158829.

